# Prediction and inference diverge in biomedicine: Simulations and real-world data

**DOI:** 10.1101/327437

**Authors:** Danilo Bzdok, Denis Engemann, Olivier Grisel, Gaël Varoquaux, Bertrand Thirion

## Abstract

In the 20^th^ century many advances in biological knowledge and evidence-based medicine were supported by p-values and accompanying methods. In the beginning 21^st^ century, ambitions towards precision medicine put a premium on detailed predictions for single individuals. The shift causes tension between traditional methods used to infer statistically significant group differences and burgeoning machine-learning tools suited to forecast an individual’s future. This comparison applies the linear model for identifying *significant* contributing variables and for finding the most *predictive* variable sets. In systematic data simulations and common medical datasets, we explored how statistical inference and pattern recognition can agree and diverge. Across analysis scenarios, even small predictive performances typically coincided with finding underlying significant statistical relationships. However, even statistically strong findings with very low p-values shed little light on their value for achieving accurate prediction in the same dataset. More complete understanding of different ways to define ‘important’ associations is a prerequisite for reproducible research findings that can serve to personalize clinical care.

---

‘Change your statistical philosophy and all of a sudden different things become important’ Steven Goodman

## Introduction

Inference and prediction are two sides of a coin when inquiring human health and disease (1–3). Let’s take diabetes mellitus as a motivating example. The inference paradigm is effective to establish biological effects that provide some insight into what leads to disturbed blood sugar levels. Diabetes in children can be a result of insufficient production of insulin hormone in the pancreas (type 1). Diabetes in adults may also reflect deficient insulin receptor response in body cells (type 2). Diabetes can moreover affect previously healthy pregnant women (gestational type). The clinical manifestation of disturbed blood sugar probably underlies partly diverging pathophysiology, which may encourage other therapeutic interventions that have been shown to yield statistically significant benefits for a particular patient group. Classical inferential statistics can also substantiate clinical observations that most patients with type 1 diabetes profit from injecting missing insulin, while obese patients with type 2 diabetes are more likely to profit from surgical intervention and symptoms in pregnant patients usually resolve after delivery.

Instead of substantiating the presence of group effects in disease biology and clinical treatment, the prediction paradigm aims to detect statistical regularities that generalize to the future. Diabetes can be diagnosed “superficially” by pattern-recognition algorithms based on frequent urination or increased thirst, possibly combined with age and gender, or its later consequences like retina damage or kidney impairment. Recognizing such symptom constellations is possible without detailed understanding of the biological processes that led to or maintain the disease. In treatment, predictive pattern-extraction algorithms can make it possible to engineer an insulin pump for accurate forecasting of the sugar response regularities that characterize a patient’s metabolism. Similar individualized predictive monitoring may enable risk prognosis and early intervention before onset of symptoms or longer-term consequences to improve medical care, without requiring understanding the metabolic mechanisms at play. In this way, inference and prediction have important but different contributions to make to biomedical research: We want to extent scientific knowledge of disease and we want to know what may happen next to an individual.

Classical inference has been intimately linked to statistical null-hypothesis testing and drawing conclusions from data guided by p-values. This framework emerged in the first half of the 20^th^ century (4, 5) for use with tools like linear regression, *t*-tests, and ANOVA. Electrical calculators not yet widely available (6, 7), this was a time when data were rare and expensive to acquire (6, 8). Hence, research experiments were often carefully designed in advance and well-controlled. The historical context also explains why classical inference was originally intended for answering research questions in subjects recruited to the local laboratory that can be addressed by transparent statistical models with few knobs to tweak (i.e., model parameters) (8, 9). Many early statistical inventions were intended to yield understanding of the relationship between a few candidate measures that were handpicked guided by the scientific question and prior research. Many of today’s medical doctors and biomedical investigators have been “raised” with this statistical culture at the university. If the scientific goal is to examine whether an effect exists or which specific input variables have most impact on an outcome, classical null-hypothesis testing remains the gold standard (10). However, a few authors, including John Ioannidis, have cast doubt that computing only p-values to draw statistical inference will continue to play an invariably important role for biomedical research (11): “With the advent of big data, statistical significance will increasingly mean very little because extremely low p-values are routinely obtained for signals that are too small to be useful even if true.”

Around the turn of the century, the rapidly increasing availability of whole-genome sequencing and high-resolution imaging ushered biomedical research into the era of “big data” (9, 12, 13). There is growing momentum for the creation, curation, and collaboration of massive datasets. For instance, the UK Biobank has gathered genetic, behavioral, environmental, and lifestyle data for extensive phenotyping of 500,000 volunteers - the currently largest biomedical data resource of its kind (www.ukbiobank.org). Due to the parallel improvements in data availability, computing power, and data storage (14, 15), the realm of data-analysis has potentially expanded faster in the last two decades than ever before (9, 12). Flexible prediction algorithms are particularly well suited for searching through rich data to extract subtle patterns (8). Such predictive modeling approaches can be less transparent but promise improved clinical translation of single-patient prediction in a fast, cost-effective, and pragmatic manner. This goal of empirically justified predictive success is sometimes viewed as a less noble science (16). Nevertheless, pioneering studies have now demonstrated the potential of “deep learning” algorithms in medicine (17) to 1) predict the cardiovascular risk, blood pressure, and smoking behavior from retina scans using medical data from almost 300,000 patients (18), 2) detect different heart arrhythmia as well as cardiologists in electrocardiograms from 30,000 patients (19), and 3) diagnose malignant skin cancer as well as dermatologists using almost 130,000 pictures (20).

There is tremendous potential in the practical goal to exploit predictive relationships for clinical endpoints in complex medical data. Needless to say, such empirical success does not fully satisfy the scientific curiosity to understand the primary biology of diseases like diabetes. Carefully planned and expensive experiments to confirm or reject a-priori verbalized research hypotheses in animals and humans will surely remain a cornerstone to generate biomedical knowledge. In this computational investigation, we therefore try to bring widespread approaches to classical inference and pattern prediction to the same table to illuminate their characteristic commonalities and differences.

## Methods

### What do we mean by ‘inference’?

The term has been used by several quantitative fields with varying, sometimes conflicting definitions (8). Here we adopt the technical meaning common in the statistical null-hypothesis testing context (21). Classical inference is aimed at scientific discovery by trying to reveal “true” properties of the studied phenomenon. Quantifying whether an effect exists in the world is especially suited to ask scientific questions like ‘Does a genetic polymorphism *contribute to* or *have an effect on* a disease?’ Providing such insight as a service to science is typically achieved by making probabilistic assumptions about how the observed data arose (e.g., the bell-shaped Gaussian distribution). The underlying structure of a scientific process is typically explored by trying to understand the way a set of input measures affect an outcome. The inference paradigm is especially useful to judge the individual relevance of each quantitative measure in impacting the response of interest. The investigator wishes to draw inference by quantitatively isolating the more important measures among the set of candidate variables, which were often hand-chosen based on existing knowledge. This intention explains why, historically, many empirical sciences have long relied on linear model approaches, even if the “true” relationship in nature is thought to be more complicated (21). Modeling for inference is self-consistent in assuming that the ‘fitted’ specified model is a sufficient summary of the studied phenomena, where each variable and its units have an immediate semantic interpretation. Often combined with careful experimental control and formally backed up by mathematical theory, the inference agenda is how traditional academic statistics has routinely dealt with small to medium datasets from planned data acquisition (8).

### What do we mean by ‘prediction’?

Describing aspects of the inner workings of the studied phenomenon is conceptually distinct from empirical research for the sake of prediction. To accurately model the world in this way, the investigator wants to extract knowledge of regularities searching through possibly meaningful candidate patterns (22, 23). This modeling goal is for instance especially suited to ask ‘Is there a set of genetic polymorphisms *useful* to *detect* whether an individual has a disease or not?’ Compared to modeling for inference, there tends to be smaller concern for the data-generating process. Prediction accuracy is the core metric to capture how well the quantitative model can *emulate* a high-level description of mechanisms in nature; that is, how well the built model can reproduce the studied phenomenon that has been quantitatively measured in the data. In the extreme case, the quantitative model may embody the discovered statistical relationship in a way that is opaque to the investigator (e.g., many “deep” neural-network algorithms). The prediction paradigm strives for highly accurate guesses by explicitly checking the fitted model by external validation. The ‘trained’ quantitative model is built for prediction in new individuals whose outcome information we would only obtain in the future. Typically, the predicted outcomes cannot be easily obtained, are expansive, or hard to come by (24). This aspect of automatically “filling in” missing information also explains why mere correlation between two variables, such as in Pearson’s correlation, may represent a more limited notion of foretelling yet-to-be measured observations (25). Out-of-sample prediction has been an important focus of activity in the more recent machine-learning community (2) and corresponds to how data analysis is often practiced in data-intensive industries (26).

### Using the linear model for inference

To assess which variables have a statistically significant relation to the outcome, we evaluated the strength of evidence using ordinary linear regression. Many statisticians have a preference for assessing significance by considering all candidate measures in the same model (cf. 27, 28), rather than carrying out simple linear regression on one independent variable at a time. A single input variable can turn out to be insignificant by itself, but become significant when part of a model with other input variables (29). The common approach to perform least-squares regression optimizes the following objective:

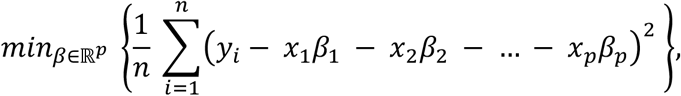

where *n* is the number of individuals who contributed to the dataset, *p* is the number of input variables *x*_*i*_ (called *independent, explanatory* or *predictor variables*) measured for each individual, and *y* is the outcome measure (called *dependent* or *explained variable*) that is to be expressed as a weighted sum of the variables *x*. The data *x* were standardized by mean centering to zero and variance scaling to one. This linear combination is estimated by fitting the *β* coefficients to all observations in the dataset. Given that the other variables are also present in the model, the approach can answer questions about the relative contributions of each of the input variables in explaining the output y. The probability model assumed that the data are sufficiently described by means and variances (21). The fitted model is assumed to encapsulate a description of how the particular input measures increased or decreased in parallel with each other to jointly explain variability in the response of interest.

After model estimation, statistical inference was drawn to decide whether the contribution of input variable *x*_*i*_ in explaining the response *y* is sufficient to be deemed *statistically significant*. The relevance of the effects is computed based on the confidence intervals of the beta coefficients (30). Inferential conclusions are drawn by formally testing for deviance of the observed effects from the null-hypothesis (e.g., a gene is not associated with diabetes) in line with the alternative hypothesis (e.g., a gene is associated with diabetes). The ensuing p-values for the input variables indicated whether our data provides enough evidence against the null hypothesis of no relevant relationship. The approach attempts to reject the null hypothesis that the beta coefficients are truly zero, that is, bear no coherent relation to the response variable. A non-significant beta coefficient suggests that the variable can be dropped from the model with little or no impact on explaining the output variable, which is however not explicitly evaluated. In typical applications of null-hypothesis testing, the p-value is computed on the entire data from all considered individuals.

### Using the linear model for prediction

For comparison with traditional linear regression, we chose a minor extension to use the linear model as a predictive pattern-learning algorithm (31). LASSO also estimates a weighted combination of the input variables, but the goal revolves around prediction. It is arguably the simplest existing method with sparsity constraint, which enforces that not all input variables are relevant in the linear model. Each variable has the same chance to be left out in the final model tuned for prediction in new observations (29). We thus identified subsets of the input variables that allow for the strongest predictive effects. Automatic variable selection was achieved by minimizing the same optimization objective augmented with a penalty term during estimation:

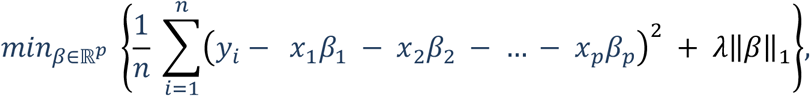

where *n* is the number of individuals who are included in the dataset, *p* is the number of input variables *x* (in this context often called *features*) measured for each individual, and *y* is the outcome to be predicted (called *target variable*) by expressing it as a weighted sum of the standardized variables *x*. The linear combination is estimated by fitting the *β* coefficients to the observations in the dataset. The hyper-parameter *λ* controls the pressure for variable selection imposed during model fitting - the degree of sparsity constraint. The higher *λ*, the stronger the tendency to set some coefficients *β*_*i*_ to exactly zero, which effectively “silences” the corresponding measure’s influence on the output variable. An explicit probability model is not required - whether the confidence intervals exceeded a threshold or not is here often no optimality criterion for variable importance. This approach did also not assume that means and variances (i.e., purely linear effects) fully describe the probabilistic mechanism in the data, only that they are informative enough to make useful predictions about the future. Once fitted, the model was applied to other samples to predict unobserved outputs or “shipped” to other laboratories for repeated application. The selected model thus automatically chose the minimal subset of predictive variables necessary for classifying for instance healthy versus diagnosed individuals. At its extreme, many pattern-learning models use the coefficient estimates as an intermediate step to achieve prediction, and actually interpreting the parameter values is little priority. In other words, many predictive modeling approaches favor the correctness of the overall prediction on new data over the individual contributions of particular beta coefficients.

Following model estimation, the practical performance of the candidate predictive model was therefore evaluated based on standard cross-validation (22). Explicit empirical guarantees are obtained to answer the question how much the predictive algorithm can be expected to generalize to data that we would see in the future. Model parameters were thus estimated on some data while the emerging model is explicitly put to the test in some independent data from unseen individuals (32): First, the linear model was built on a larger part of the dataset. Second, emerging candidate algorithms were evaluated and selected on unused data to avoid an overly optimistic evaluation of goodness-of-fit (22). Because all conditions for independent, identically distributed observations are usually met for the left-out data, the out-of-sample prediction performance on the testing data samples can quantify how likely the same pattern could be detected in future, not yet seen patients. In this way, the cross-validation scheme quantified the *out-of-sample performance* as an estimate of a model’s capacity to generalize to data samples acquired in the future. As the LASSO does not provide a full least-squares fit due to its shrinkage property, we computed debiased out-of-sample predictions using ordinary least-squares on the collection of active variables. This common modification allowed disentangling the influence of shrinking and variable selection in forming predictions with LASSO. As an important consequence, all prediction scores reported in this work were obtained from ordinary linear regression (without shrinkage bias) based on the full set or subset of input variables automatically selected from the preceding LASSO estimation.

Such modeling for prediction, routinely practiced in many applications of pattern-recognition algorithms, is centered around evaluating the capacity of already extracted models to derive quantities of interest from new, potentially later encountered individuals. This form of building models from data has been explicitly optimized for and is naturally applicable to a single data point, such as one whole-brain scan or one sequenced genome of a particular individual. Please appreciate that it is not adviced to compute the usual p-values on the automatically selected input variables (33, 34). This is because variable selection by the LASSO is itself a random process that is ignored by the theoretical guarantees of classical inference for statistical significance (35). Put differently, data-driven model selection is undermining hypothesis-driven statistical inference. The initial prediction-based filtering step alters the sampling distribution of the variable coefficient estimates for subsequent significance-based variable filtering. This incompatibility between statistical inference and variable selection invalidates classical null-hypothesis testing and optimistically biases computed p-values (35), which is an active area of research (33, 36, 37).

### Simulations

It has been noted that formal guarantees for the expected model prediction performance are challenging to derive by mathematical theory (8, 32). In such settings, empirical simulation can come to the rescue for studying the properties of statistical methods in computational experiments (30). Here we directly confronted linear modeling for inference and for prediction in a series of synthesized datasets, columns of input variables *X*, each related or not related to the outcome *y*. Each dataset was generated from a set-up ground-truth model *y* = *βX* + *ϵ*, where *β* are fixed random coefficients, *X* is a data matrix containing *n* samples and p variables with random entries drawn from a standard Gaussian distribution 𝒩(_*μ*=0,*σ*=1_), and *ϵ* denotes the added Gaussian noise. Each dataset was fed into the linear model with the aim to identify significant input measures or to identify input measures most useful for accurate predictions on new observations (cf. above).

To sharpen the distinction between explanatory and predictive modeling in general, we systematically varied distinct aspects of the data-generating process:

i. <u>Samples-to-variables ratio:</u> To investigate the relation between the number of samples *n* relative to the number of variables *p*, we systematically varied the number of available observations. We covered the lower range between 50 and 100 samples in steps of 10, which probably well reflects a majority of studies in biomedicine. Between 100 and 2,000 samples we increased the sample size in steps of 100. Moreover, we considered the extreme cases 10,000 and 100,000 samples, which acknowledges recent large-scale datasets such as the UK Biobank. The total number of input variables was kept constant to preclude secondary effects on the results due to changing model capacity.
ii. <u>Proportion of informative variables:</u> To study how the fraction of relevant versus irrelevant variables modulate the inferential and predictive processes, we varied the proportion of non-zero *β* coefficients in the ground-truth model used for generating *X*. We considered 14 proportions ranging from only 1 to all 40 input variables carrying information about the response *y*.
iii. <u>Redundant versus unique sources of information:</u> To elucidate how correlated input measures trade-off against each other with respect to the outcome, we introduced different degrees of pairwise covariation between the variable columns of *X* (i.e., collinearity). Ground-truth models also generated data from a multivariate Gaussian distribution that exposed 50% or 90% percent of common variation between the relevant variables, complementing datasets that contain only mutually independent variables (i.e., 0% covariation).
iv. <u>Signal-to-noise ratio:</u> To assess the role of nuisance variation in the data, such as induced by imperfect measurement techniques, we systematically manipulated the noise *ϵ* in how the real model relates to the response *y*. The nuisance term was generated from 𝒩(_*μ*=0,*σ*=1_) and multiplied by 0.5, 1, 2, 5, 10, or 0 (i.e., generating data without any noise).
v. <u>Model violations:</u> To examine more closely how inference and prediction behave when the linear model can not fully capture how the data came about, we introduced pathological alterations on 50% of the relevant variables in *X*. In addition to datasets with exclusively linear effects (i.e., we can find the true model), deviations between the generating and fitting model were introduced by one of several data transformations: taking the absolute value, the natural logarithm, the exponential, the square root, the multiplicative inverse, or polynomial expansion of degree 2-5.

The collection of simulated datasets realized 113,400 different data-analysis scenarios. For each case, we focused on the best (smallest) p-value among all input variables in the model and the highest prediction performance of the overall model as quantified by the (out-of-sample) R^2^ score. All simulations were carried out on a parallel computing server with 48 Intel Xeon CPUs (1,200 - 2,900 GHz) and 62 GB of working memory. The analyses required almost 4 weeks of computation time and produced 2 GB of modeling results.

### Scientific computing implementation

Python was selected as the scientific computing engine. Capitalizing on its open-source ecosystem helps enhance replicability, reusability, and provenance tracking. The *statsmodels* package was used to estimate ordinary least squares regression and corresponding p-values (http://statsmodels.github.io). The *scikit-learn* package (38) provided efficient, unit-tested implementations for handling state-of-the-art machine-learning procedures (http://scikit-learn.org). All analysis scripts that reproduce the results of the present study are readily accessible and open for reuse (http://github.com/banilo/https://github.com/banilo/infvspred2018). The repository also provides extended Jupyter notebooks with additional analyses and an interactive WebApp.

## Results

### Simulated datasets

Across 113,400 constructed datasets (Fig. 1), we made several observations about the characteristic differences between seeking statistical inference and maximizing model prediction. Fitting linear models to series of datasets generated with increasing non-linear effects easily reached significance but distinctly varied in the predictability of the outcomes (Fig. 2F; Fig. 3). It was expected that even, as opposed to odd, polynomial data transformation (e.g., x^2^ or x^4^) incur larger violations to model validity because the direction of effects in the input variables is lost. As such, 4^th^-order polynomial expansion deteriorated model fit more than 5^th^-order expansion, entailing both worse p-values and worse R^2^ prediction performance (out-of-sample). To emulate random variation such as from measurement error, we added gradually increased noise in the data. This additional challenge during model fitting decreased the predictability more systematically than the significance (Fig. 2D). Adding more random noise to the data was not observed to entail less models with statistically significant variables. To emulate the frequently encountered challenges when facing collinear data, we have increased the correlation shared between the input measures (Fig. 2C). More variation common to several input variables appeared to worsen the p-values more than the prediction performance. Covariance of 90% yielded p-values (i.e., smallest in the model) closer to the typical p < 0.05 threshold and seldom very low p-values. Concurrently, many data-analysis scenarios that did not yield a single significant relation between an input variable and the response of interest were generated in this high-collinearity setting. To capture some implications of the ongoing trend to data aggregation in biomedicine, we gradually increased the available data points per generated dataset (Fig. 2A). At the highest sample size of n=100,000, low significance tended to more systematically agree with low predictability and extremely high significance also mostly concurred with perfect out-of-sample performance. That is, in datasets, bigger than is currently the norm, we observed more consistent correspondence between significance and prediction. Exploring different proportions of relevant measurements in the ground-truth model (Fig. 2B), we noted that fewer truly relevant inputs gave rise to strongly significant p-values in the presence of poor predictive performance. Finally, applying linear models that deviate from the data-generating process of the input and output variables (Fig. 2E) led to results with high significance and predictability in many cases. However, using the valid (linear) model to fit the randomly generated (linear) data allowed for many of the best prediction performances (Fig. 3A).

**Figure 1.**
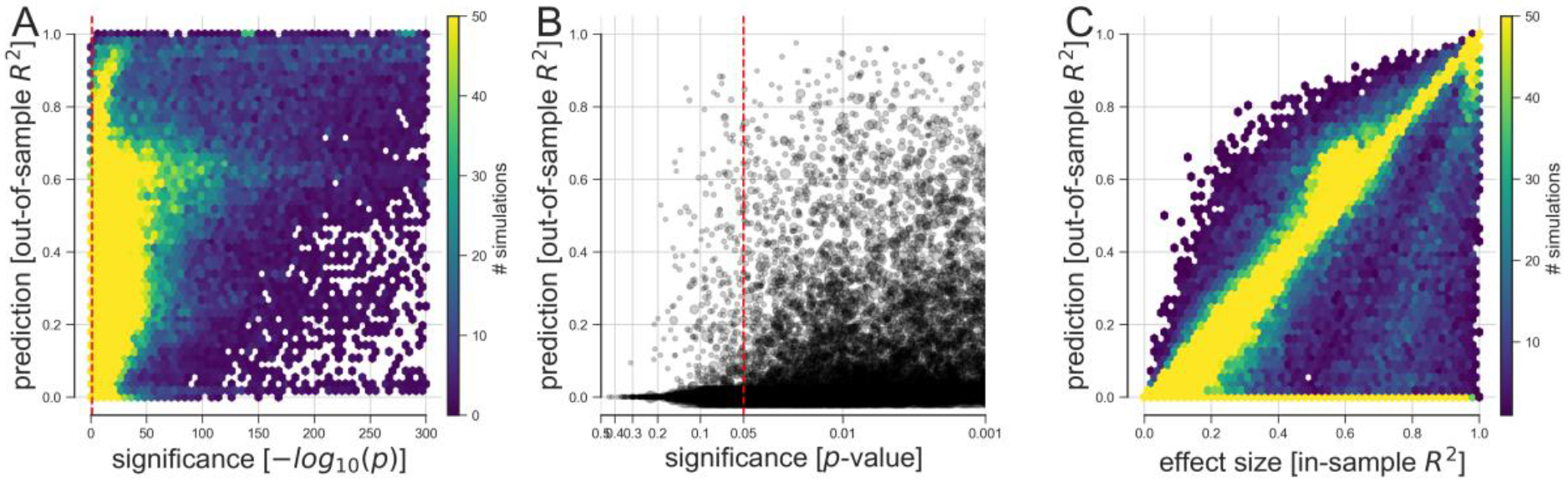
Predictability versus significance of effects in simulated datasets. Based on 113,400 simulations, the discrepancy between predictive and explanatory modeling was quantified in a wide range of possible data-analysis cases. The generated variables and outcomes were analyzed by linear models with the goal to draw classical inference (smallest p-value among all model coefficients, x axis) and to evaluate model forecasting performance on never seen data (out-of-sample R^2^ score of the model, y axis). **A)** Hexagonal binning summarizes how many simulations led to a particular relation between prediction and inference in a 2D histogram. This area-by-area visualization was proposed for aggregating data with many observations (51). **B)** Predictive accuracy and statistical significance are juxtaposed with their relation to the commonly applied thresholds at p < 0.05, p < 0.01, and p < 0.001 (bigger grey circle means bigger sample size). **C)** Prediction accuracy is compared to the effect size derived from the explained variance on the model fitting data (in-sample R^2^ score of the model). In the large majority of conducted data analyses, at least one input variable was significantly related to the response variable at p < 0.05 (red dashed vertical line). However, based on the same data, we observed considerable dispersion in how well such significant linear models were able to make useful predictions on fresh observations.

**Figure 2.**
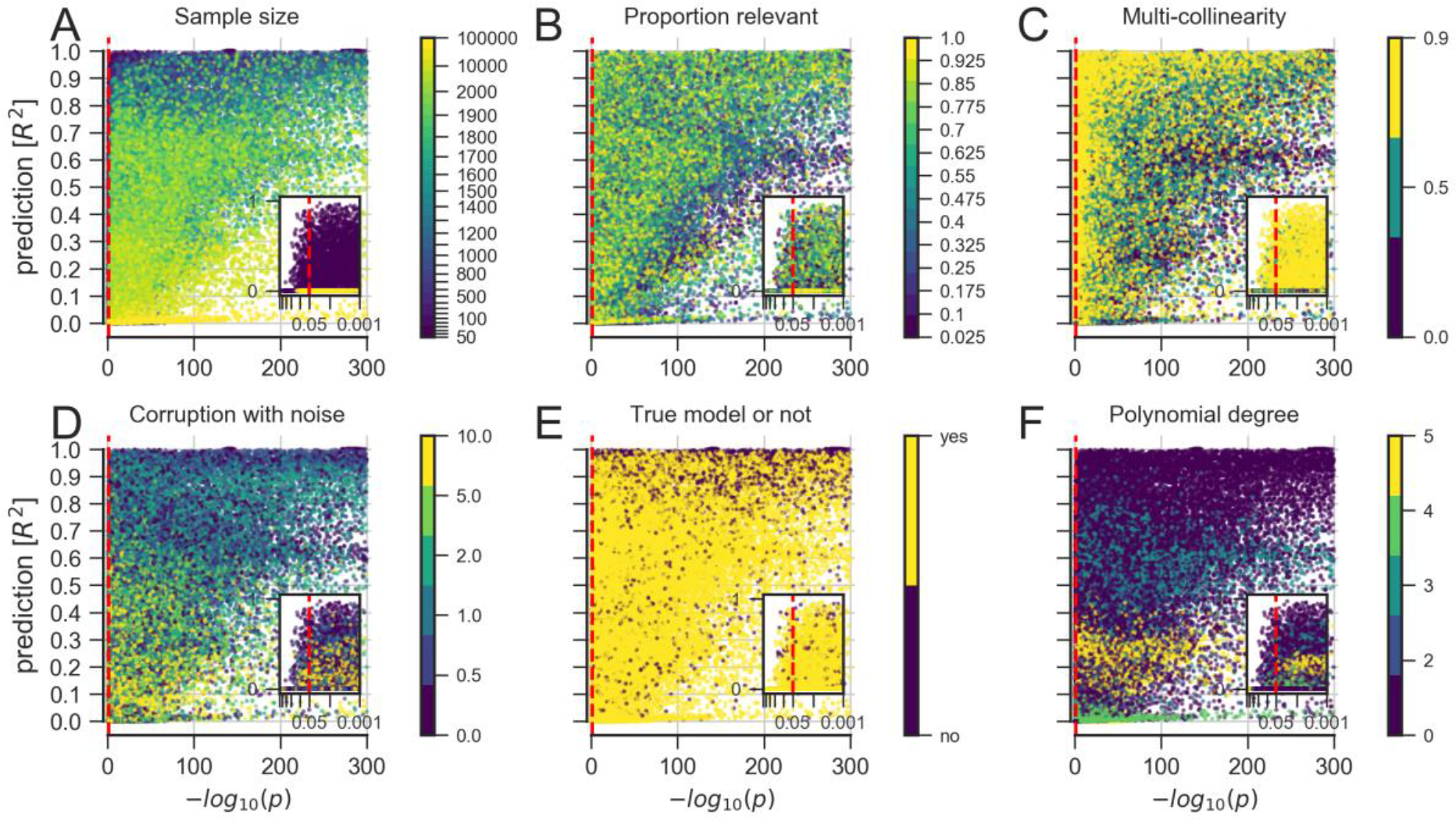
Properties underlying analysis results of simulated data. Explore in more detail how linear modeling for significance testing (smallest p-value, x axis) and linear modeling for prediction (out-of-sample R^2^ score, y axis) agreed and diverged across constructed datasets. **A)** Increasing the number of available data points eventually yielded co-occurrences of strong significance and prediction. **B)** Small numbers of relevant predictors allowed for scenarios with highly significant p-values in combination with poor predictive performance. **C)** Increasing correlation between the input measures, common in biological data, appeared to worsen the p-values more than the prediction performance. **D)** Increasing random variation in the data, which can be viewed as imitating measurement errors, appeared to decrease the predictability more systematically than the significance. **E)** Pathological settings, where the chosen model does not correspond to the data-generating process of the input and output variables, tended to enhance both significance and predictions. **F)** Fitting a linear model to data with increasing non-linear effects easily reached significance but distinctly varied in predictability of outcomes.

**Figure 3.**
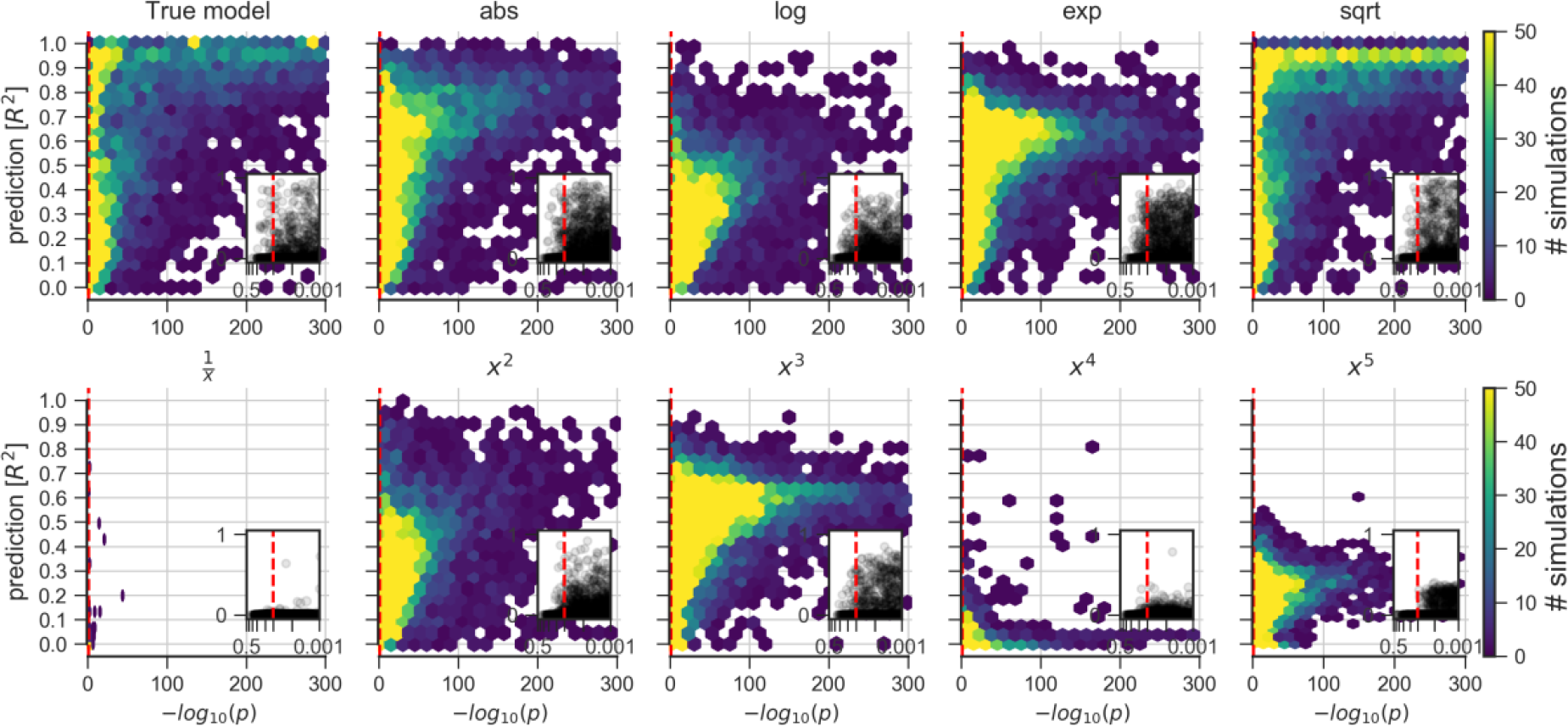
Implications of different model violations in simulated data. Explores consequences of applying a linear model to datasets that are known to contain non-linear data mechanisms of different types and degrees (cf. Fig. 2A). Certain non-linear effects are likely to influence measurements of various real biological systems. That is, in everyday data analysis, some misalignment between the data and the commonly employed linear model is likely to be the rule rather than the exception.

### Real medical datasets

To complement the simulated datasets, the same direct comparison between explanatory modeling and predictive modeling was carried out in common real-world datasets (Fig. 4). The quantitative re-evaluation is presented here for four medical datasets that are frequently used as examples in data-analysis teaching and textbooks (e.g., 22, 29).

**Figure 4.**
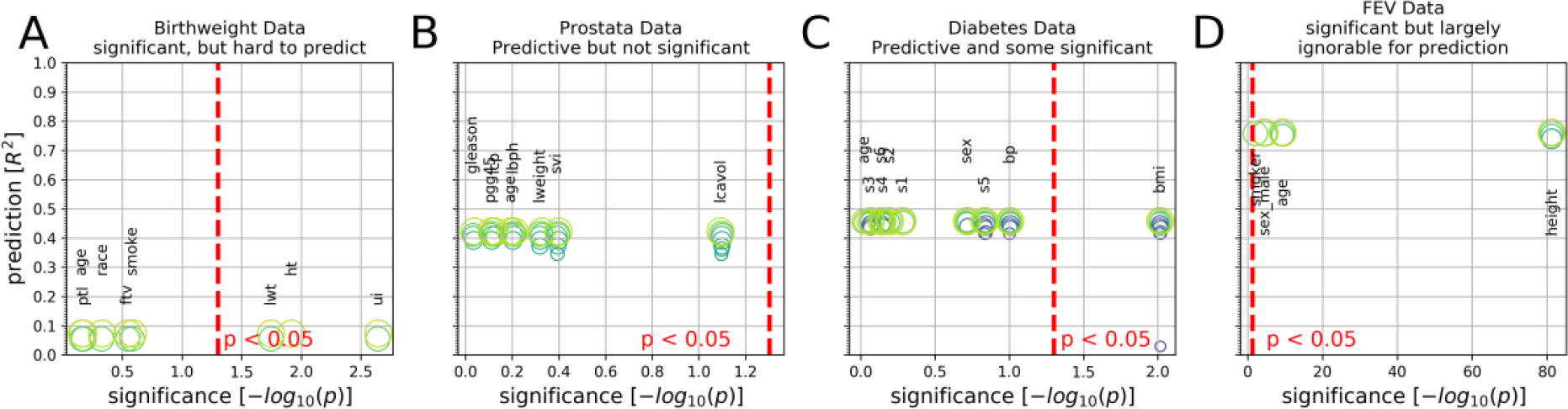
Predictability versus significance in four medical datasets. Integrative plots summarize the inferential importance of each linear model coefficients (p-values on *x-axis*, log-transformed) and the predictive importance of coefficient sets (out-of-sample R^2^ scores on *y-axis*, obtained from model application on data not used for model fitting). **A)** The body weight is to be derived from 8 measures in 189 newborns. 3 out of 8 measures are statistically significantly associated with birth weight at p < 05 *(red line)*. Yet, using the linear model for prediction explained only 8% of the variance in new babies (R^2^=0.08). **B)** Prostate specific antigen (PSA), a molecule for prostate carcinoma screening, is to be derived from 8 measures in 87 men. None of the 8 coefficients reached statistical significance based on common linear regression, although the fitted coefficients of the predictive model achieved 42% explained variance in unseen men. **C)** Disease progression after one year is to be derived from 10 measures in 442 diabetes patients. Body mass index (BMI) gave the only significant coefficient (p=0.01), which alone however explained only an estimated 3% of disease progression in future patients. The full coefficients of the predictive model achieve 46% explained variance in independent patients. **D)** Lung capacity as quantified by forced expiratory volume (FEV) is to be derived from 4 measures in 654 healthy individuals. All measures easily exceeded the statistical significance threshold. However, a predictive model incorporating body height alone performed virtually on par with predictions based on all 4 coefficients (R^2^=0.74 versus R^2^=0.76). In sum, linear models can show all combinations of predictive vs. not and significant vs. not in biomedical data analysis.

In the <u>birthweight dataset,</u> standard linear regression was used to evaluate the relation of 8 candidate measures to the body weight of 189 newborn babies (Fig. 4A). In this approach, the 3 effects that reached statistical significance at p < 0.05 comprised the mother’s weight at the last menstrual period (p=0.018, lwt), existing history of hypertension (p=0.012, ht), and presence of uterine irritability (p=0.002, ui). The in-sample model fit amounted to R^2^=0.141. In the prediction setting, linear models were trained and evaluated involving the same data. The best estimate of the explained variance expected in babies that we would see in the future reached only R^2^=0.08 (as measured by unbiased out-of-sample prediction) based on the full set of 8 input measures. After predictive variable selection “silenced” the influence of the age of the mother and the number of physician visits during the first trimester (ftv), the remaining 6 active measures still allowed for a prediction performance of R^2^=0.06. These appeared to be a predictive core subset among the input measures because at 5 out of 8 coefficients the linear model prediction diminished to be worse than the average model. Comparing the strongest measures identified by classical inference and pattern prediction by explicit model checking on the birthweight data, a few input variables easily reached significance. However, relying on the same data, it was challenging to obtain a predictive model with convincing pattern generalization to new data points, despite the reasonable sample size.

In the <u>prostate cancer dataset,</u> none of 8 input measures turned out to be statistically significantly associated with prostate-specific antigen (PSA) in 87 men (Fig. 4B). This molecule is widely used by medical doctors for cancer screening and monitoring to guide whether or not to surgically remove the prostate gland. Cancer volume (lcavol) was closest to being judged important with p=0.081. In contrast, the estimated prediction accuracy achieved R^2^=0.42 with 8/8 coefficients, R^2^=0.42 with 5/8 coefficients, R^2^=0.38 with 3/8 coefficients, and still R^2^=0.35 with 2/8 coefficients. Notably, the single most useful measure to predict a man’s PSA concentration in these data was the cancer volume with an explained population variance of R^2^=0.25 with 1/8 coefficients (lcavol). That is, despite lacking statistical significance, we found coherent predictive patterns in the data that were reliably extracted. The combined information from several variables was required to achieve the higher prediction performances. The prediction approach also detailed that lcavol > svi > lweight carry the most relevant information to forecast a man’s PSA level. The ordered ranking coincided with the absolute beta coefficients obtained using linear regression. In the prostate cancer dataset, in-sample model estimation reverberated with (all three positive) variable importance in out-of-sample prediction performance but was in disagreement with the obtained insignificant p-values.

In the <u>diabetes dataset</u>, disease progression after one year was to be derived from 10 measures in 442 patients (Fig. 4C). In modeling for inference, only the body mass index (bmi) was deemed significant at p=0.01 among all input variables. This single measure, however, only accounted for 3% of explained disease progression in the population when modeling for prediction. Adding another predictive variable - s5 - to the linear model with bmi, enhanced the prediction accuracy to R^2^=0.42. Adding more and ultimately all input variables into the model led to small additional improvements in prediction performance (R^2^=0.46). In fact, s5 showed the highest positive beta coefficient (at the beginning of the regularization path, where small sparsity constraint was imposed) but did not turn out as the final variable remaining in the model. Summing up the results on the diabetes data, the single significant variable carries negligible information to achieve reliable prediction in new data; only when s5 is incorporated in the predictive model, very good predictions was achieved in new patients not yet witnessed by the model.

Finally, in the <u>FEV dataset</u>, the lung capacity captured as forced expiratory volume (FEV) was to be derived from 4 measures in 654 healthy individuals (Fig. 4D). All input variables easily reached the statistical significance threshold. Yet, a predictive model built from the same data revealed that considering body height alone performed virtually on par with predictions based on all 4 coefficients (R^2^=0.74 versus R^2^=0.76). That is, age, gender and smoking habits all easily reached statistical significance, but offered little added value for the purpose of prediction. In the case of lung capacity prediction, the predictive variable selection concurred with the highest absolute coefficient in both approaches to determined importance. Here the prediction regime has probably missed the mechanistically relevant influence of smoking on lung capacity by pragmatic predictions based on body height alone. The high significance of all input variables may have been facilitated by the comparably high sample sizes.

## Discussion

Exploring a battery of empirical simulations and several biomedical datasets offered insight into asymmetric tendencies between seeking accurate predictions in new individuals and identifying statistically significant effects across individuals. While prediction and inference share a first step of estimating linear-model coefficients, the difference arises in what the data analyst decides to do next with the fitted model.

Charting a broad spectrum of data analysis scenarios possible in everyday research, statistically significant relationships were not always a guarantee to also achieve successful predictions when applying the model to other individuals. To restate, effects robust at the conventional significance level of p < 0.05 varied between virtually no and almost 100% explained variance in fresh data. By contrast, effects not significant at p < 0.05 mostly failed to deliver useful predictions in data from unseen individuals. In short, predictability appears to be a demanding criterion because even small predictive performances typically coincided with finding underlying significant statistical relationships in almost all cases. However, even statistically strong associations with very low p-values shed only modest light on their value for the goal of prediction based on the same data.

Researchers in most empirical sciences face questions of data analysis. What does it mean that a measure is ‘important’ or not? Statistical significance identified important variables based on (in-sample) deviation from a theoretical non-effect that is unlikely explained by noise. Out-of-sample prediction, instead, discarded unimportant variables if the omission did not diminish the empirical model performance on unseen data. P-values were computed by whether an input measure would take the actually obtained value at most 1 in 20 times if its impact on the outcome is not important. An official report of the American Statistical Association (ASA) emphasized that ‘Statistical significance is not equivalent to scientific, human, or economic significance’ (10). Hence, an association between a candidate gene and diabetes grounded in a statistically significant p-value may not necessarily imply that the same gene can be used to successfully predict whether a given individual is affected by that disease. On a related note, in psychology and other empirical sciences (39–42), there is accumulating evidence for a replication crisis. Significant results published in a scientific paper are in many cases not substantiated when the identical experiments and data analyses are conducted again at a later point in time. We used a predictive method considered variable ‘importance’ in a different way. A variable was considered relevant if leaving it out hurt the overall prediction accuracy when applying the previously built model was explicitly checked on fresh observations (2). Some authors believe that such empirical validation procedures to establish importance may increase in the future due to adoption of code and data sharing. These expanding practices can promote across-study and across-method confirmation (43).

In fact, ‘importance’ in quantitative research has probably no uniform theoretical basis (2, 44); and can therefore take different forms and shapes even in the canonical linear model. A statistical method that produces importance assessments still requires the informed judgment of the investigator how far the conclusions should be trusted. The initial choice of analysis method may be more or less well aligned with the substantive research question. Put differently, using p-values or prediction accuracies for backing up research claims have both flaws and each is insufficient in some way (1, 27, 29). The ASA statement recommended: ‘No single index should substitute for scientific reasoning’ (10) - a viewpoint shared by other prominent investigators (45, 46). In particular, Ioannidis and colleagues recently stressed monocultural training of biomedical scientists in statistical null-hypothesis testing as one reason behind frequent misuses of statistical methods (47).

### Conclusion

Our quantitative investigation exposed how linear models - a workhorse in many areas of biomedical research - can be used with distinct and partly incompatible motivations. Using these tools for the purpose of inference is ideal to uncover characteristics of biological processes. Using linear modeling for the alternative purpose of prediction is particularly suited for pragmatic forecasting of biological processes, potentially including clinical endpoints in individual patients. Some statisticians therefore proposed that data-analysis applications should be primarily distinguished by the modeling goal, rather than strictly cataloguing each method under an umbrella term, such as ‘statistics’ versus ‘machine learning’, ‘hypothesis-based’ versus ‘data-driven’, or ‘confirmatory’ versus ‘exploratory’ (43, 48). It is critical for investigators and practicing medical doctors to acknowledge the incongruent modeling philosophies of drawing statistical inference and seeking algorithmic prediction, as well as their non-identical scopes of interpretation (2, 49). Statistical literacy may become increasingly relevant for taking rigorous and reproducible steps on our journey to personalied medical care, with the prospect to benefit the well-being of suffering patients.

More broadly, the prediction-inference dilemma may also remind us of some ideas of Claude Bernard - a pioneer of controlled experiments in biomedicine (50). Prediction may be closer to what he called ‘empirical medicine’ oriented towards practical patient care as an often theory-free endeavor, such as symptom monitoring, risk assessment, and choosing therapeutic intervention. Statistical inference may bear a more direct relationship to his conceptualization of ‘scientific medicine’ aimed at elucidating unknown principles underlying biological processes driven by theory, such as asking for the reasons why certain individuals are at risk for disease onset or illuminating why a certain drug works better in some individuals than others.

In approaching a future of precision medicine, it may become central that mainstream statistics or machine learning are related but importantly different. We demonstrated that diverging conclusions can emerge even when the data are the same and widespread linear models are used (8). Awareness of the strengths and weaknesses of both “data-analysis cultures” is unavoidable to fully benefit from the accelerating data deluge in biology and medicine.

## Acknowledgements

We thank several investigators for insightful comments on a previous version of the manuscript: B.T. Thomas Yeo (National University of Singapore), Guillaume Dumas (Institut Pasteur/France), Nikolaus Kriegeskorte (Columbia University/USA), Daniele Marinazzo (Ghent University/Belgium), Benjamin de Haas (University of Giessen/Germany), and João Sato (Universidade Federal do ABC/Brazil).

DB was funded by the Deutsche Forschungsgemeinschaft (DFG, BZ2/2-1, BZ2/3-1, and BZ2/4-1; International Research Training Group IRTG2150), Amazon AWS Research Grant (2016 and 2017), the German National Merit Foundation, as well as the START-Program of the Faculty of Medicine (126/16) and Exploratory Research Space (OPSF449), RWTH Aachen. DE acknowledges support by the Amazon AWS Research Grant (2015), the German National Merit Foundation, as well as the French National Institute for Informatics and Automation (INRIA) (Starting Researcher Position SRP 2016). The authors declare no competing interests.

